# A cell wall-associated gene network shapes leaf boundary domains

**DOI:** 10.1101/2021.11.15.468678

**Authors:** Nathalie Bouré, Alexis Peaucelle, Magali Goussot, Bernard Adroher, Ludivine Soubigou-Taconnat, Eric Biot, Zakia Tariq, Marie-Laure Martin-Magniette, Patrick Laufs, Nicolas Arnaud

## Abstract

Boundary domains delimit and organize organ growth throughout plant development almost relentlessly building plant architecture and morphogenesis. Boundary domains display reduced growth and orchestrate development of adjacent tissues in a non-cell autonomous manner. How these two functions are achieved remains elusive despite the identification of several boundary-specific genes. Here, we show using morphometrics at the organ and cellular levels that leaf boundary domain development requires SPINDLY (SPY), an O-fucosyltransferase, to act as cell growth repressor. Further we show that SPY acts redundantly with the CUP-SHAPED COTYLEDON transcription factors (CUC2 and CUC3), which are major determinants of boundaries development. Accordingly at the molecular level, CUC2 and SPY repress a common set of genes involved in cell wall loosening providing a molecular framework for the growth repression associated with boundary domains. Atomic force microscopy (AFM) confirmed that young leaf boundary domain cells have stiffer cell walls than marginal outgrowth. This differential cell wall stiffness was reduced in *spy* mutant. Taken together our data reveal a concealed *CUC2* cell wall associated gene network linking tissue patterning with cell growth and mechanics.

**Summary statement:** Decreased cell-wall loosening gene expression contributes to the coordination of cell growth and mechanics with tissue patterning thus driving boundary development.

## Introduction

Boundaries act both as frontiers to separate adjacent tissues or organs and as organizing centers providing positional clues to control the fate of neighboring cells (Dahmann et al., 2011; Irvine and Rauskolb, 2001). Thus boundary domains are required to correctly pattern developing organs. For instance in animals, defects in boundaries lead to developmental abnormalities including impaired wing or brain development (Dahmann et al., 2011). In contrast with the determinate development occurring in animals, plants continuously form new aerial growth axes separated from the shoot apical meristem to build their architecture. These new growth axes can either produce new branches or give rise to specialized lateral organs such as leaves or flowers. Independently of their fate, all lateral organs are separated from the meristem by boundary domains, which delimitate cell territories and orchestrate their development (Aida and Tasaka, 2006). Despite decades of efforts to decipher their functions, plant boundary domains remain an elusive population of cells for which little information is available.

The patterning and maintenance of boundary domains rely on the activity of the CUP-SHAPED COTYLEDON transcription factors (CUC) which belong to the NAC transcription factor family (Aida et al., 1997; Vroemen et al., 2003). There are three *CUC* genes in Arabidopsis *CUC1, 2* and *3. CUC1* and *CUC*2 mRNA but not *CUC3* are targeted by a *miRNA, MIR164* (Laufs et al., 2004). The CUC transcription factors regulate both shoot meristem formation (Aida et al., 1999) and correct organ separation in various developmental contexts (Aida et al., 1997; Burian et al., 2015; Gonçalves et al., 2015). Accordingly, CUC2 and CUC3 are key regulators of leaf shape through their roles on leaf margin development (Blein et al., 2008; Hasson et al., 2011; Nikovics et al., 2006), *CUC1* being not expressed during leaf development (Nikovics et al., 2006). During leaf development, CUC2 and CUC3 define boundary domains at the leaf margin - called sinuses - allowing differential growth to shape the leaf. At the cellular level, the coordinated activity of CUC2/3 transcription factors locally suppress growth and have a positive effect at a distance on the initiation and maintenance of high growth rate probably *via* a mechanism involving auxin (Bilsborough et al., 2011). The CUC2 transcription factor acts through the activation of *CUC3* and *KLU/CYP78A5*, encoding a Cytochrome P450, which serves as molecular relays through the modulation of auxin signaling pathway (Maugarny-Calès et al., 2019). Acting downstream of *CUC2, CUC3* maintains reduced growth of the boundary domains *via* the control of cell growth through unknown molecular mechanisms (Serra and Perrot-rechenmann, 2020). Accordingly, *cuc2* loss-of-function mutants fail to initiate teeth while in *cuc3* loss-of-function mutant teeth growth is not maintained (Hasson et al., 2011). Several hormonal pathways impinge on boundary domains establishment (Hepworth and Pautot, 2015). For instance Brassinosteroid (BR) have been shown to antagonize boundary domains formation through the down regulation of *CUC* genes (Gendron et al., 2012). Low BR levels are maintained within boundary domains by the activation of BAS1, a cytochrome P450 involved in BR catabolism, thus leading to the reduced growth of boundary domains (Bell et al., 2012). Auxin also plays a fundamental role during boundary domains establishment, nicely exemplified by its implication to leaf serration development. Other regulatory molecules have recently emerged as important regulators of boundary domains. The EPF/EPFL secreted peptides and the ERECTA family receptors contribute to boundary domains formation both during leaf development and ovule initiation probably through modulation of auxin responses (Kawamoto et al., 2020; Kosentka et al., 2019; Tameshige et al., 2016).

In an attempt to identify new actors of boundary domains, we previously performed a genetic suppressor screen of a line over-expressing *CUC2* and identified *MUR1*, coding for a GDP-D-mannose 4,6-dehydratase involved in GDP-L-fucose production. More specifically, we showed that L-Fucose contributes to boundary domain establishment in various developmental contexts (Gonçalves et al., 2017). Fucose is a hexose incorporated to xyloglucans, rhamnogalacturonan II and arabinogalactans in plant cell wall (O’Neill et al., 2001; Van Hengel and Roberts, 2002) and added to protein through the activity of specific fucosyltransferases (Strasser, 2016). Therefore *mur1* mutants, which are deficient in GDP-L-Fucose, present pleiotropic developmental phenotypes such as small plant stature and reduced leaf development (O’Neill et al., 2001). In order to understand in which pathway L-Fucose is involved to regulate boundary domains establishment, we analyzed leaf serration formation as a proxy for boundary development in mutants lacking specific fucosylation events. Using a loss-of-function mutant for MUR2, a fucosyltransferase specifically targeting xyloglucans, and a transgenic line overexpressing the xyloglucan fucosyl hydrolase AXY8/FUC95A both displaying non-fucosylated xyloglucans (Günl et al., 2011; Vanzin et al., 2002), we showed that xyloglucan fucosylation was not necessary for leaf serration development (Gonçalves et al., 2018).

Recently SPINDLY (SPY) has been described as a O-fucosyltransferase able to target REPRESSOR OF GA (RGA), a negative regulator of the gibberellin signaling pathway from the DELLA family (Zentella et al., 2017), as well as PSEUDO-RESPONSE REGULATOR 5 (PRR5), a core circadian clock component (Wang et al., 2020). *spy* loss-of-function mutants have been originally identified in a genetic screen for plantlet resistant to the GA biosynthesis inhibitor Paclobutrazol (Jacobsen and Olszewski, 1993). This mutant displays constitutive GA phenotypes suggesting that SPY negatively regulates GA signaling pathway (Jacobsen et al., 1996). The confirmation that SPY may directly O-fucosylates RGA provides a molecular framework for the function of SPY in the GA-signaling pathway. However, as *spy* mutants do not completely resemble WT plants treated with GA (Swain et al., 2001), it is likely that SPY acts as well through GA-independent pathways. This has been recently shown during root development where SPY regulates root hair patterning in a GA-independent pathway (Mutanwad et al., 2020). Accordingly, SPY has been proposed to positively regulate Cytokinin (CK) signaling, highlighting a central role in the regulation of GA/CK crosstalk throughout plant development (Greenboim-Wainberg et al., 2005). This GA/CK hormonal crosstalk is instrumental to maintain *KNOX* (Class I *KNOTTED1*-like homeobox)-dependent meristematic activity (Hay et al., 2002; Jasinski et al., 2005), which implies that SPY has a crucial role during SAM development and/or maintenance. Additionally to this function in the SAM, several reports show that *spy* mutants have altered leaf development with little or no serrations at their margins (Greenboim-Wainberg et al., 2005; Maymon et al., 2009) but detailed analysis of the implication of SPY in these boundary domains development is lacking.

Here, we precise the implication of SPY to leaf development and more generally to boundary domain development. While early serration initiation is not altered in the *spy-3* mutant, the growth maintenance of the formed tooth is affected. The analysis of cell parameters reveals that SPY is required to maintain restricted growth of the sinus cells thus mimicking a *CUC3* loss-of-function mutant. Further, our genetic analysis suggests that *SPY* acts redundantly with *CUC2* and *CUC3* to control boundary domains development. Comparing transcriptomic data of CUC2-regulated genes with available SPY-regulated genes reinforce our genetic observations. Indeed both SPY and CUC2 regulate a common set of genes controlling cell wall properties. Accordingly, we show that CUC2 represses cell growth independently of CUC3 possibly *via* modifications of cell wall mechanics. Together this work provides a molecular framework for the role of SPY during boundary domain development where SPY and CUC2 act through a common pathway and reveals a concealed growth repressive function for CUC2 involving cell expansion.

## Results

### *SPY* regulates leaf morphogenesis

Several *spy* mutant alleles (Greenboim-Wainberg et al., 2005; Maymon et al., 2009; Steiner et al., 2012) have been reported to display leaves with little or no serrations but this leaf phenotype was never fully characterized. Therefore, we quantified the leaf shape of the loss-of-function *spy-3* mutant. The *spy-3* mutant contains a G>A transition changing a Glycine to a Serine residue at the position 593 of the SPY protein, resulting in a non-functional protein (Jacobsen et al., 1996) (Fig S1). *spy-3* mutant and WT mature leaves displayed similar blade lengths while their widths hence their blade area were smaller in *spy-3* (Fig S2A, B and C). Both morphometrics and dissection index (DI) calculations - a global descriptor of leaf complexity (Gonçalves et al., 2017) - show that *spy-3* leaves are smoother than WT leaves (Fig 1A,B,C). To go further we measured the shape of the second tooth - as the ratio between tooth high and tooth width - in this dataset and show that WT serrations are pointer than *spy-3* serrations (Fig 1D). Two additional alleles namely *spy-22* and *spy-23* (Fig S1 and Fig S3) had smoother leaves that the WT with less pronounced serrations, ruling out *spy-3* specific bias on leaf morphology. Furthermore, a *spy-22* mutant expressing *pSPY::SPY-FLAG* construct had a restored WT leaf shape phenotype (Fig S3). Taken together our results show that *SPY* is involved in the development of leaf serrations.

**Figure 1.**
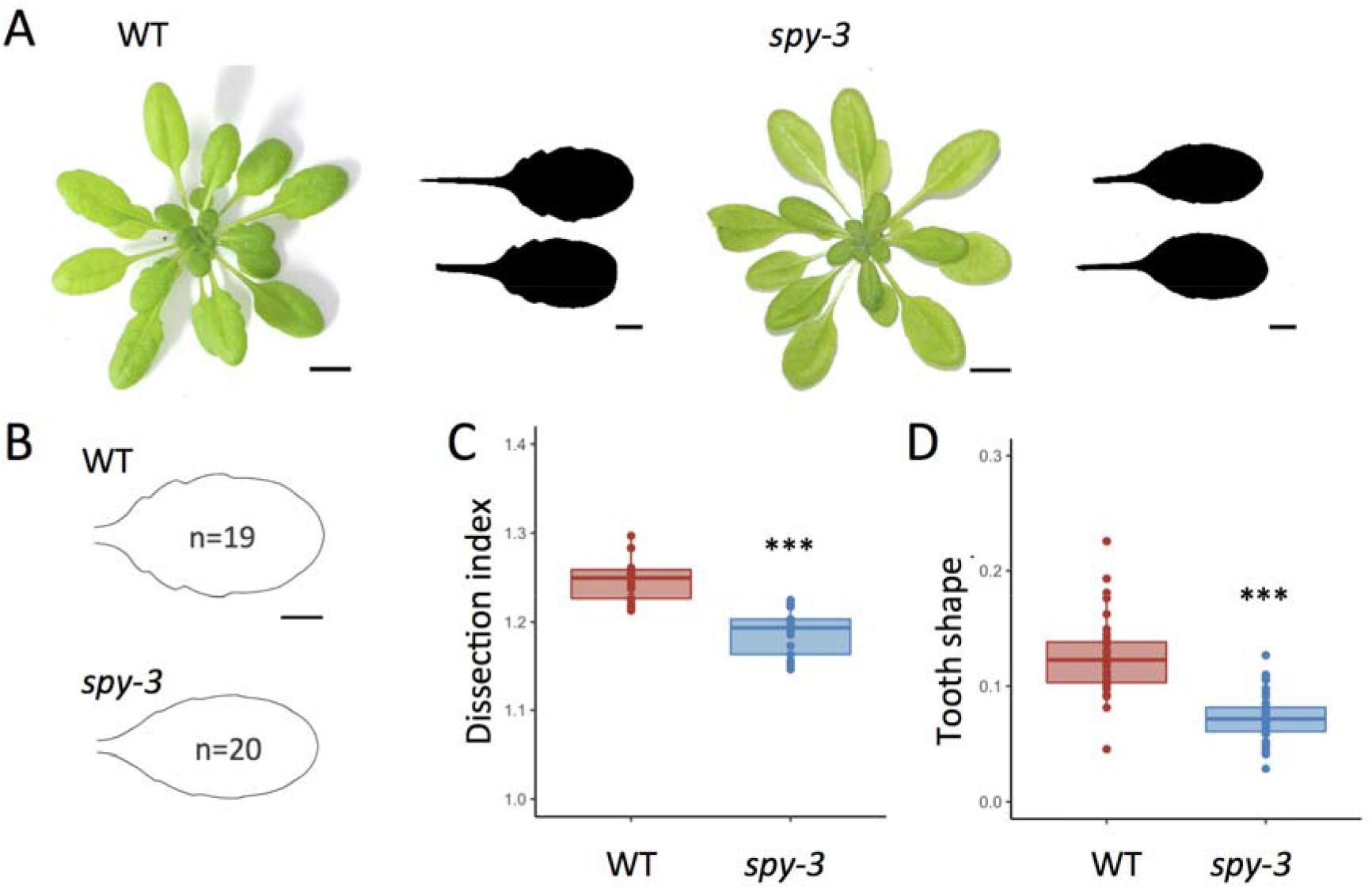
Morphometric analysis of *spy-3* mature leaf shape. A. Wild type and *spy-3* mutant rosette from plants grown in short-day conditions for 6 weeks. Representative silhouettes from mature leaves from ranks 11-12-13 are also represented. Scale bar = 1cm. B. Mean shape of mature leaves of wild type and *spy-3* mutant. Scale bar = 1cm. C. Quantification of *alpha-hull* normalized dissection index (DI) for WT and *spy-3* mature leaves. D. Quantification of the shape of the second tooth from WT and *spy-3* mature leaves. B,C,D. WT (n=19) and spy-3 (n=20) 6-week-old leaves grown in short-day conditions ranks 11-12-13. Statistical significance (Student’s test) is designated by *** p<0.0001.

### SPY is required for growth serration maintenance

As mature leaf shape results from the sum of processes occurring at different developmental times, it is important to access growth kinetic data of the *spindly* mutant to conclude about SPY precise roles during leaf shape development. To do so, we reconstructed detailed developmental trajectories of *spy-3* loss-of-function mutant leaves using the *Morpholeaf* software from a set of leaves from ranks 11, 12 and 13. As both the leaf initiation rate and the leaf growth rates are comparable between *spy-3* and the WT up to 6 mm length (Fig. S4), we chose to limit our morphometric analysis to this early stage and use leaf blade length as a proxy for leaf developmental stage. Tooth 1 height is drastically reduced in *spy-3* from early stages and never reaches WT values (Fig 2A) while tooth1 width is not modified at early developmental stages (up to 3mm) (Fig 2B), resulting in sharper teeth in the WT (Fig 2C). Together our data suggest that SPY is involved in teeth growth maintenance. As maintenance of tooth growth is associated with the definition of the leaf boundary domain at the sinus (Hasson et al., 2011; Maugarny-Calès et al., 2019), we analyzed sinus angle as a local parameter related with the local growth repression at the sinus. Although the evolution of the sinus angle for the distal sinus of tooth 1 throughout its development has comparable dynamics both in *spy-3* than in WT, *spy-3* sinus angle is always less pronounced than WT sinus angle (Fig 2D). These data suggest that the alteration of leaf shape may partially result from local defects in boundary domain definition in the *spy-3* mutant.

**Figure 2.**
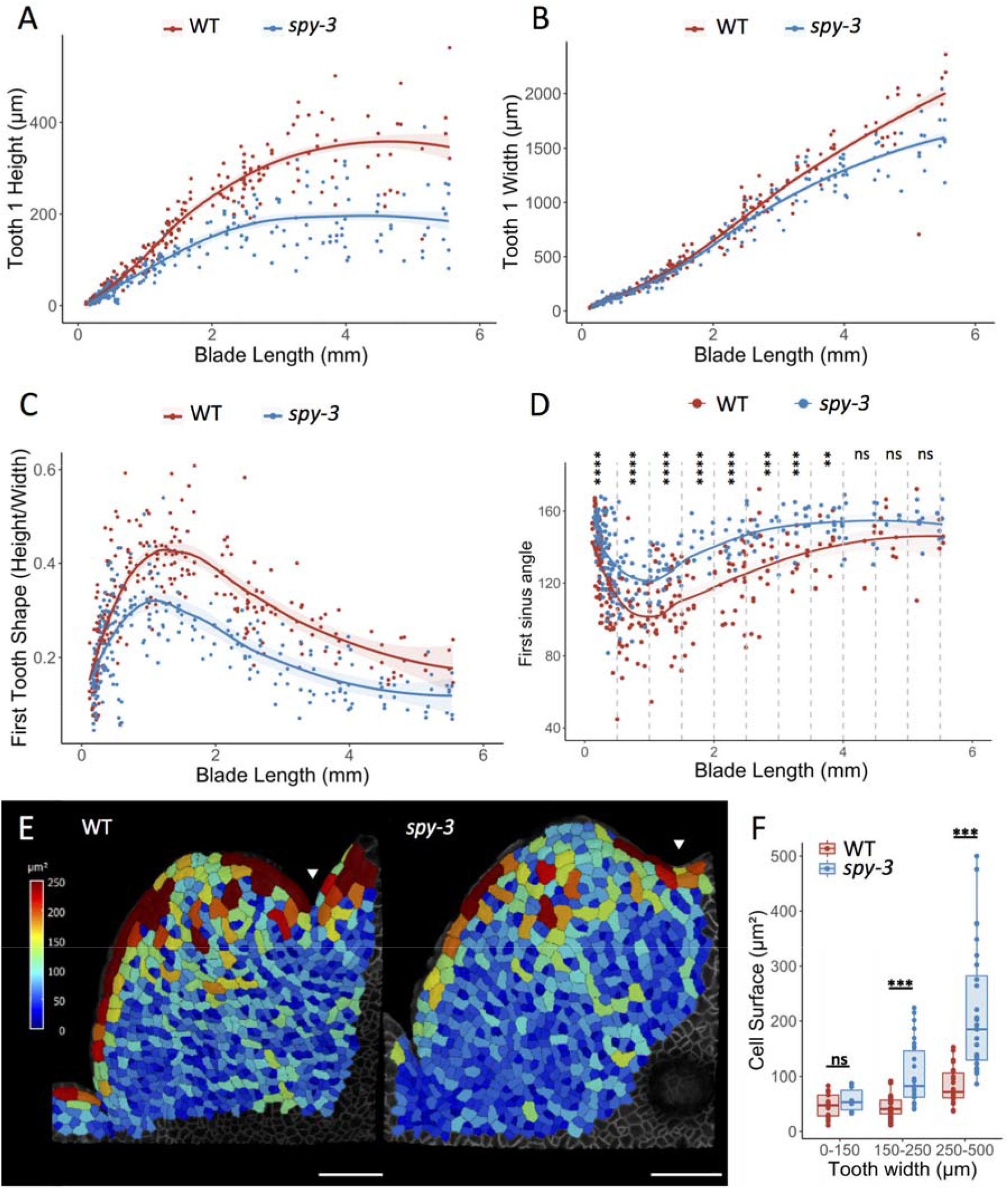
Developmental kinetics and cell size quantification during *spy-3* serration development. A. First tooth height plotted against blade length for WT and *spy-3*. B. First tooth width plotted against blade length for WT and *spy-3*. C. Tooth shape of the first tooth, calculated as tooth height over tooth width plotted against blade for WT and *spy-3*. A,B,C. Leaf ranked 11-12-13 dissected throughout their development from WT (n=190) and *spy-3* (n=194) plants grown in short-day conditions were used. Each tooth is represented by a dot, and a LOESS curve is shown for visual interpretation. D. Mean first sinus angle measured in short-day grown WT (n=190) and *spy-3* (n=194) and plotted against blade length. 250 μm-wide classes were made to perform statistical analysis. Statistical significance (Student’s test) is designated by ** p<0.01, *** p<0.001, **** p<0.0001. E. Representative cell area heatmaps for WT and *spy-3* from the first tooth sinus cells. Arrowheads indicate the crease defining the first apical sinus on each tooth. Scale bars = 50 μm. F. Projected surfaces quantification from the first tooth sinus cells plotted against tooth width. WT (n = 17), *spy-3* (n = 6) for [0-150] μm. WT (n = 37), *spy-3* (n = 30) for in [150-250] μm. WT (n = 22), *spy-3* (n = 28) for in [250-500] μm. Statistical significance (Student’s test) is designated by ns = not significant, *** p<0.001.

### SPY is required to inhibit sinus cell growth during leaf development

During leaf development, spatial differences in cell growth rate sustain tooth outgrowths (Serra and Perrot-rechenmann, 2020) which are integrated at the leaf level leading to final leaf shape. As sinus angle was altered in the *spy-3* mutant compared to the WT, we set out to analyze sinus at the cellular level. The distribution of cell surface in 3D acquisitions for tooth 1 of leaves from ranks 11, 12 and 13 was hence measured in both genotypes (Fig 2E). We focus on the first distal sinus to limit bias due to the mechanical constraints of the previous tooth outgrowth. The shape of the tooth and the depth of the sinus were different between *spy-3* and the WT in these 3D acquisitions, which confirmed the data from the 2D developmental kinetics. In order to specifically assess the size of the sinus cells, we analyze the Gaussian curvature of the 3D projected surface. Sinuses were identified as surface areas that exhibit a negative Gaussian curvature (Serra and Perrot-rechenmann, 2020) and the surfaces of sinus-specific cells were measured from independent leaves. Sinus cell surfaces were then plotted according to tooth width for both *spy-3* and WT leaves (Fig 2F). For teeth 1 up to 150 μm wide, sinus cell sizes of early leaf primordia were not significantly different between *spy-3* and the WT. Later for teeth ranging from 150 to 250 μm and 250 to 500 μm, sinuses of *spy-3* mutant leaves are constituted of larger cells than WT cells. Our data show that SPY is required to maintain restricted sinus cell growth at late stages of tooth development. Interestingly, the *cuc3-105* loss-of-function mutant has been described to have bigger cells at the sinus due to local release of cell growth (Serra and Perrot-rechenmann, 2020). Thus *spy-3* and *cuc3-105* mutants display very similar sinus cell phenotypes suggesting that SPY and CUC transcription factors have similar roles for sinus cell development probing the question whether they function through a common pathway to coordinate growth restriction of sinus cells.

### SPY, CUC2 and CUC3 act redundantly during boundary domains development

To check whether SPY acts in a *CUC*-dependent pathway to define boundaries, we first analyzed the genetic interaction with *CUC2g-m4*, a mutated version of *CUC2* with altered *MIR164*-target site, leading to a local over-expression of *CUC2* mRNA and to very serrated leaves (Maugarny-Calès et al., 2019; Nikovics et al., 2006). *CUC3* which is acting downstream of *CUC2* has already been shown to reduce leaf serration of the *CUC2g-m4* line (Hasson et al., 2011). CUC2 over-expression leaf phenotypes (measured by DI calculations) were suppressed in *spy-3* or *cuc3-105* background (Fig 3A). Furthermore we used the *mir164a* loss-of-function mutant as an alternative way of increasing *CUC2* levels. Consistently, the over-serrated leaf phenotype of *mir164a* was suppressed by the *spy-3* mutation (Fig 3A). Together these genetic data suggest that CUC2 requires SPY or CUC3 activity to control leaf serration development.

**Figure 3.**
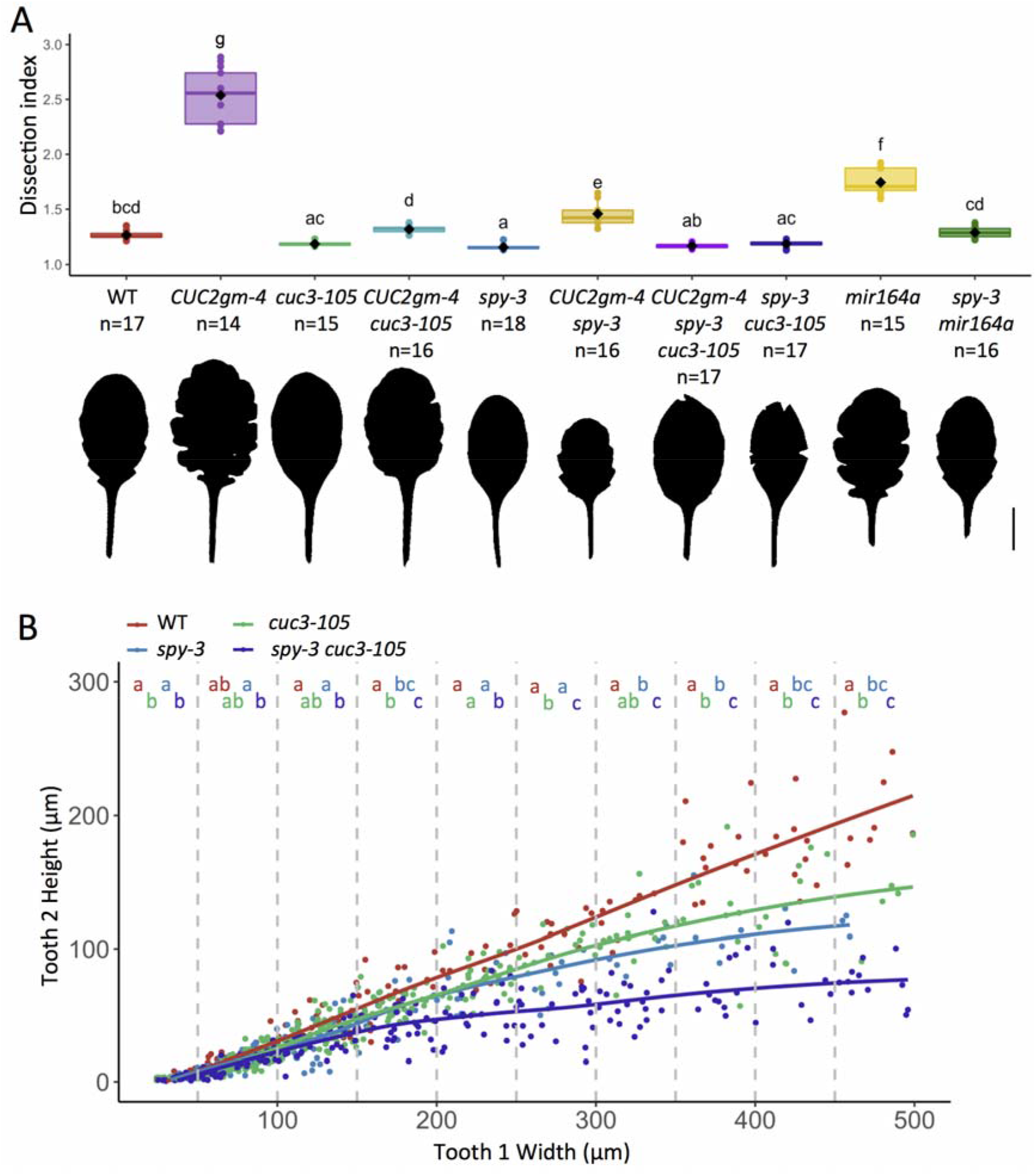
*CUC* and *SPY* genetic interactions during leaf development. A. Quantification of *alpha-hull* normalized dissection index (DI) for WT (n = 17), *CUC2g-m4* (n = 14), *cuc3-105* (n = 15), *CUC2g-m4 cuc3-105* (n = 16), *spy-3* (n = 18), *CUC2g-m4 spy-3* (n = 16), *CUC2g-m4 spy-3 cuc3-105* (n = 17), *spy-3 cuc3-105* (n = 17), *mir164a* (n = 15) and *spy-3 mir164a* (n = 16) mature leaves of ranks 11-12-13. A representative leaf silhouette is shown for each genotype analyzed. Scale bar = 1cm. B. Analysis of *CUC3* and *SPY* genetic interactions during leaf development. First tooth height from WT (n = 190), *spy-3* (n = 194), *cuc3-105* (n = 213), *spy-3 cuc3-105* (n = 191) plotted against 50μm-wide first tooth width classes. Short-day grown plants from 22 to 50 days after sowing (DAS) were used. Statistical analysis: one-way ANOVA analysis was performed within each class, followed by Tukey comparison test.

As *spy3* and *cuc3-105* mutants have very similar phenotypes both at the organ and at the cellular scales, and that they both suppress CUC2-highly serrated leaves phenotype, we hypothesize that *CUC3* and *SPY* act in the same genetic pathway to control serration development. To test the idea, we generated a double *spy3 cuc3-105* mutant and analyzed its leaf developmental trajectory (Fig 3B). As CUC3 maintains tooth growth by locally inhibiting the growth of sinus cells, we used tooth height of the first tooth (T1) as a proxy of CUC3 activity. Strikingly, when tooth height of T1 was measured for different tooth width classes, *cuc3-105* and *spy-3* display comparable quantitative phenotypes while the two mutations together have an additive effect on tooth height suggesting that they act independently on leaf shape (Fig 3B). Accordingly, *spy-3* and *cuc3-105* mutations have also an additive effect on the over-serrated *CUC2gm-4* phenotype (Fig 3A). These data imply that alternative routes exist to restrict growth at the sinus independently of *CUC3*. In order to decipher the relative contribution of *SPY, CUC2* and *CUC3* to boundary cell growth, we decided to analyze their roles during cotyledon rather than during leaf development because no serrations are initiated when CUC2 activity is altered. Both double mutant *spy-3 cuc3-105* and *spy-3 cuc2-1* show stronger cotyledon fusion phenotypes compared with the corresponding simple mutants showing that SPY acts redundantly with CUC2 and CUC3 to define boundaries (Table 1 and Fig S7). This result shows also that SPY acts in different developmental contexts and suggests that SPY contributes more generally to the definition of developmental boundary domains.

**Table 1.** Quantification of cotyledon fusion defects in *cuc2-1, cuc3-105* and *spy-3* mutants combinations.

| Genotype | Phenotype |  |  |  | Total seedlings |
| --- | --- | --- | --- | --- | --- |
|  | normal (%) | weak (%) | mild (%) | strong (%) | n |
| *col* | 99,37 | 0 | 0,63 | 0 | 315 |
| *cuc3-105* | 99,52 | 0 | 0 | 0,48 | 631 |
| *spy-3* | 100 | 0 | 0 | 0 | 317 |
| *spy-3 cuc3-105* | 82,79 | 4,66 | 10,39 | 2,17 | 645 |
| *cuc2-1* | 100 | 0 | 0 | 0 | 320 |
| *spy-3 cuc2-1* | 82,57 | 6,59 | 10,52 | 0,31 | 637 |

### *SPY* and *CUC2* act through a common molecular network to restrict sinus cell growth

Our genetic analysis suggests that *CUC2, CUC3* and *SPY* redundantly restrict boundary domains growth. As we saw a local growth defect in the *spy-3* mutant, we first tested whether *SPY* was expressed together with the *CUC* genes within the leaf boundary domains. We used *pSPY::SPY-GFP* reporter line crossed with either *pCUC3::CFP* or *pCUC::RFP* to monitor simultaneously *SPY* localization and *CUC* gene expression patterns. Although SPY is broadly expressed in leaf epidermis at early developmental stages, it overlapped with *CUC2* and *CUC3* within leaf boundary domain cells (Fig S5). In addition, we show that even though *CUC2* levels vary grandly between *cuc2-1* mutant, the WT and the *CUC2gm-4* line, the expression levels of *SPY* do not change suggesting that SPY expression is not regulated by CUC2 (Fig S6).

As our genetic analysis shows that CUC2 activity requires *SPY*, we next wondered how this translates at the molecular level. Previous transcriptomic analysis identified genes differentially expressed in *spy-3* compared to WT (Qin et al., 2020). To identify CUC2 downstream elements we performed a transcriptomic profiling of an activated CUC2 DEX-inducible line. Among the differentially expressed genes, we found that about 20% of the genes up-regulated in the *spy-3* mutant were down-regulated upon CUC2 induction (datasetS1). Indeed, we identified 2569 genes down-regulated 6 hours after *CUC2*-induction (FDR<0.05) and among them, 100 are up-regulated in the *spy-3* mutant which represents a significant proportion of the 494 genes up-regulated in total in *spy-3* (Hypergeometric test, *p-value* = 6.06E-14) (Fig 4A). Gene ontology analysis performed using this set of 100 genes reveals an enrichment in genes related to plant-type cell wall (GO:0009505, enrichment 9.65, raw *p-value* = 4.31E-05 (Fisher exact test), FDR = 4.45E-02) with a function related to cell wall organization and biogenesis (GO:0071554, enrichment 7.38, raw *p-value* = 1.13E-06 (Fisher exact test), FDR = 6.75E-03). Among these genes, we identified several genes coding for xyloglucan endotransglucosylase/hydrolases (XTH4, XTH15, XTH18 and XTH19), arabinogalactan proteins (AGP4, AGP7, AGP9 and AGP12), as well as two genes coding for expansin-like (EXLA1 and EXLA2). These genes contribute to the cell wall loosening as relaxing enzymes. Indeed XTH18 and XTH19 were both previously shown to be involved in the control of hypocotyl growth, as overexpressing lines for *XTH18* and *XTH19* both promoted hypocotyl growth in the dark (Miedes et al., 2013). In addition, AGP4, AGP7, AGP9 and AGP12 were identified in a large-scale gene expression pattern study on fast-growing seedlings as robust markers of growth (Kohnen et al., 2016). Similarly to what was observed for XTH proteins, an *EXLA2* overexpression is able to increase growth in dark-grown hypocotyls (Boron et al., 2015). In addition, a biomechanical analysis of the *EXLA2* overexpressing line showed that the cell wall resistance was decreased in the hypocotyl, suggesting that EXLA2 may modify the cell wall organization and composition (Boron et al., 2015). Therefore our transcriptomic analysis suggests that cell wall remodeling enzymes are the functional elements acting downstream of CUC2 and SPY. Importantly, we have independently validated our transcriptomic data as we found that both *EXLA1* and *EXLA2* transcript levels are reduced after *CUC2* induction in another set of experiments (Fig 4B).

**Figure 4.**
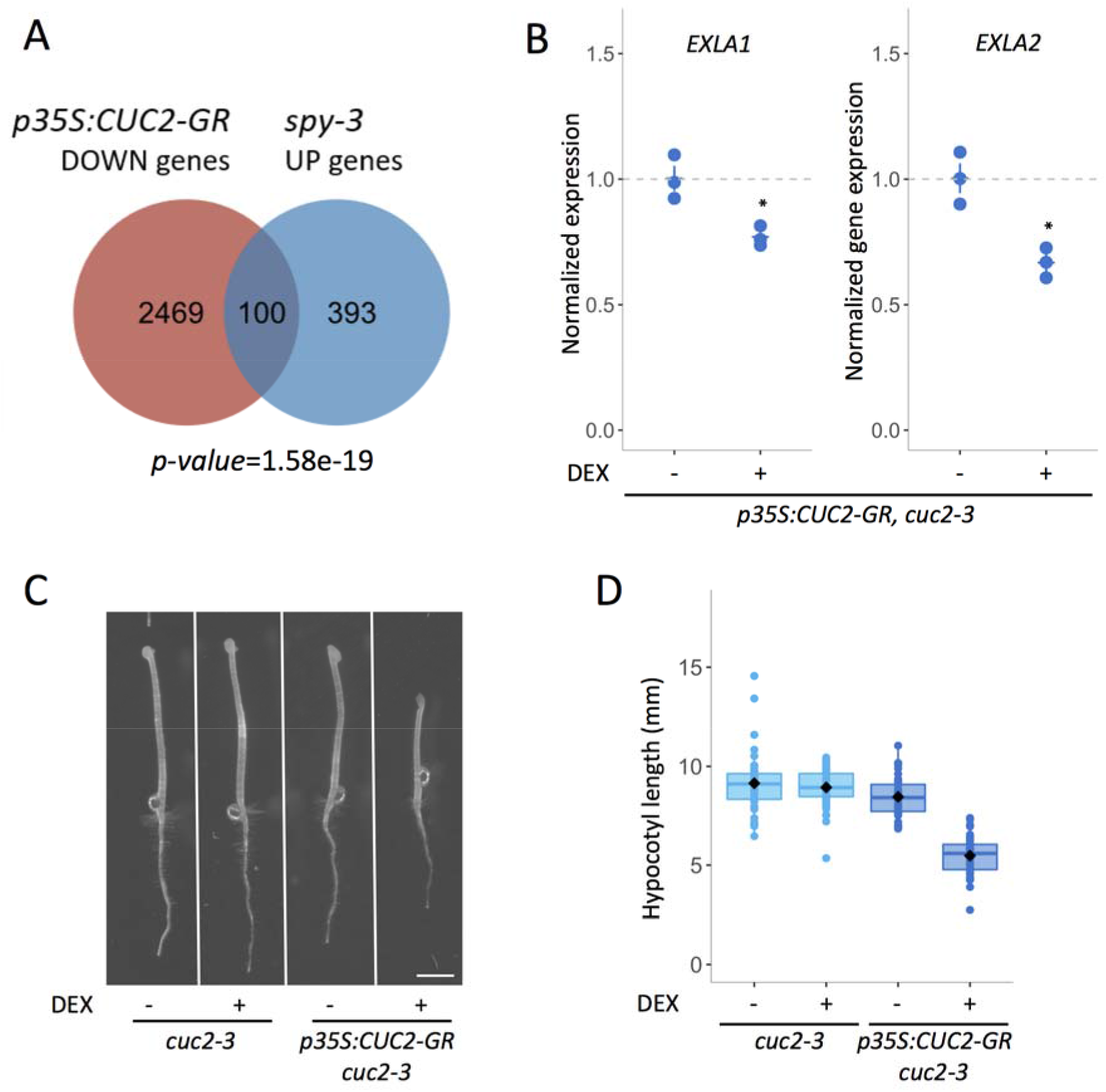
CUC2 inhibition of dark-induced hypocotyl elongation is associated with down-regulation of cell wall relaxing genes. A. A total of 2,569 genes were down-regulated in DEX-induced *p35S:CUC2-GR* line. 493 genes up-regulated in *spy-3* mutant compared with WT were identified. Numbers below Venn diagrams correspond to hypergeometric probability (N_total *At* genes_ = 33,602) (over-enrichment based on the cumulative distribution function (CDF) of the hypergeometric distribution). B. Expression level of *EXLA1* and *EXLA2* in a *35S:CUC2-GR cuc2-3* line dark-grown for 72 hours *in vitro*, treated either with mock treatment or with 10 μM DEX. Each dot represents a biological RNA sample. *EXLA1* and *EXLA2* transcript levels were measured by real-time quantitative RT-PCR normalized by *EF1*α and *Actin2*. Statistical significance (Student’s test) is designated by ns = not significant, * p<0.05. C, D. Representative phenotypes (C) and corresponding hypocotyl length quantification (D) in *35S:CUC2-GR cuc2-3* line and *cuc2-3* control dark-grown for 72 hours *in vitro*, treated either with mock treatment or with 10μM DEX.

To test whether *CUC2* overexpression may indeed inhibit cell expansion, we analyzed the effect of CUC2 induction upon DEX treatment during the elongation of dark grown hypocotyl which results mostly from the cell elongation rather than cell division (Gendreau et al., 1997). Dark grown hypocotyls are significantly shorter when CUC2 is induced with smaller hypocotyl epidermal cells compared with non-induced conditions (Fig 4B and 4C). These results are coherent with the reduced levels of expression of genes linked with cell wall plasticity after CUC2 induction. Taken together, our data suggest that the overexpression of the CUC2 transcription factor is sufficient to repress a set of genes which function relate to cell wall loosening providing a plausible molecular framework for the activity of CUC2.

### CUC2 represses cell elongation independently of CUC3

The activity of CUC2 is mediated by CUC3 which acts as a molecular relay (Maugarny-Calès et al., 2019). As CUC2 and CUC3 redundantly control boundary domain development, it is possible that CUC2 alters cell elongation independently of CUC3. As no serrations are initiated in a *cuc2* loss-of-function mutant, this hypothesis has been difficult to test during leaf development. Here, we have the opportunity to test whether the CUC2-dependent growth repression function that we have highlighted with the DEX-induced CUC2 line in dark-grown hypocotyl depends on CUC3. When CUC2 is DEX-induced in absence of CUC3, dark-grown hypocotyls are shorter with smaller epidermal cells than in non-induced conditions (Fig 5A and 5B) showing that CUC2 acts independently of CUC3 on cell elongation in the dark-grown hypocotyl model. In order to check whether this reduction of growth is associated with changes in cell wall related gene expression, we monitored their mRNA accumulation upon CUC2 induction in dark grown hypocotyls. In a *cuc2-3 cuc3-105* mutant background, CUC2 induction is sufficient to drastically reduce the accumulation of *XTH18, XTH19, EXLA1* and *EXLA2* mRNA showing that CUC2 inhibits their expression independently of *CUC3* (Fig 5C). These results suggest that changes in cell wall gene expression triggered by CUC2 may counteract hypocotyl cells elongation in the dark.

**Figure 5.**
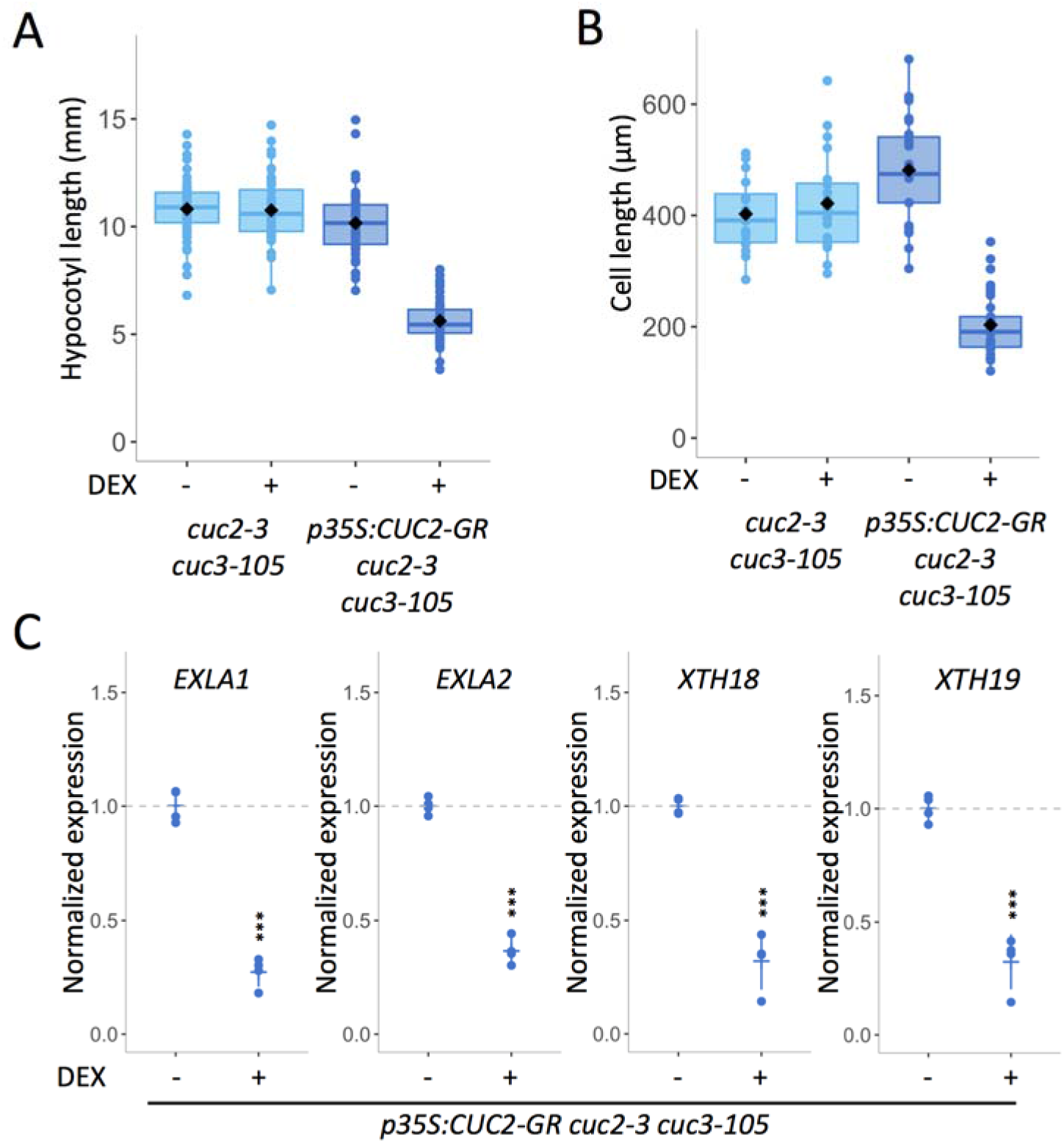
CUC2 inhibits dark-induced cell elongation independently of CUC3. A, B. Hypocotyl length quantification (A) and hypocotyl cell length quantification (B) in *35S:CUC2-GR cuc2-3 cuc3-105* line and *cuc2-3 cuc3-105* control dark-grown for 72 hours *in vitro*, treated either with mock treatment or with 10 μM DEX. C. Expression level of *EXLA1, EXLA2, XTH18 and XTH19* in a *35S:CUC2-GR cuc2-3* line dark-grown for 72 hours *in vitro*, treated either with mock treatment or with 10 μM DEX. Each dot represents a biological RNA sample. Transcript levels were measured by real-time quantitative RT-PCR normalized by *EF1*α and *Actin2*. Statistical significance (Student’s test) is designated by ns=not significant, *** p<0.001.

### Cell wall mechanics at the leaf margin

Our molecular data support a role for CUC2 in the control of cell wall properties. In order to check whether this is also the case in the organs where CUC2 is expressed, we used Atomic Force Microscopy (AFM) to measure the cell wall stiffness of sinus cells-where *CUC2* is expressed - and compare with the cell wall stiffness of tooth cells - where *CUC2* is not expressed. In young leaf primordia, when the tooth starts to emerge as shown by topological images obtained using AFM microscopy, cells of the margins do not display different sizes between sinus and tooth domains in WT (Fig 6A and Fig 6B). Yet, sinus and tooth show a differential relative stiffness: cell sinus walls being stiffer than the cell tooth walls (Fig 6A-B-E). This is consistent with both the expression pattern of *CUC2* (Woodson and Chory, 2008) and our molecular data showing that CUC2 inhibits the expression of genes known to promote cell wall loosening. To validate these data, we quantified the Apparent Young’s modulus (*Ea*) to evaluate the elasticity of the leaf margin tissue in another set of experiment analyzing cell wall stiffness in 12 young dissected leaves for the WT. This shows once again that sinuses domains are consistently stiffer than tooth domains (Fig 6F). This result provides a mechanical framework for the development of boundary domains.

**Figure 6.**
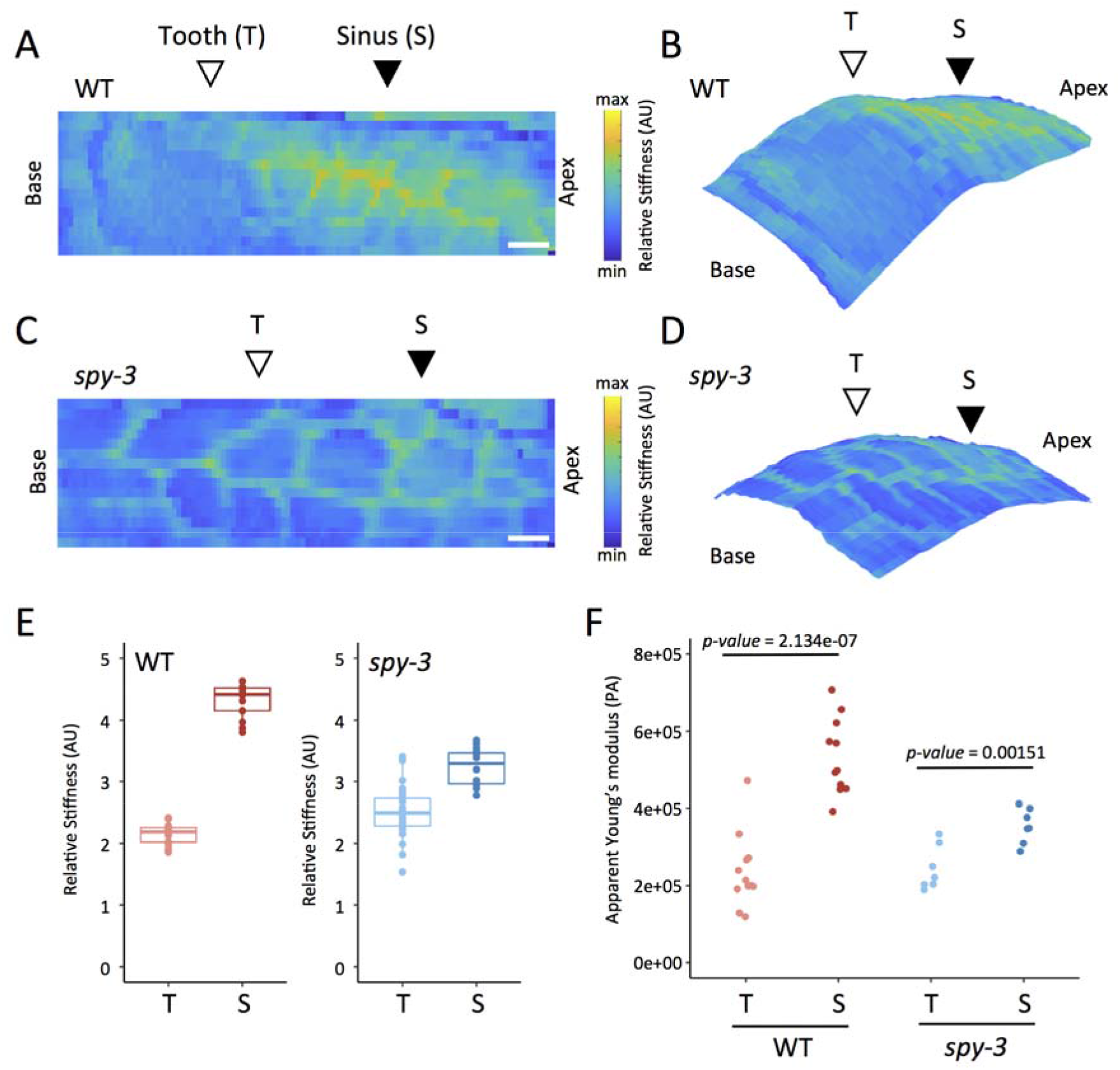
Mechanics at the leaf margin of WT and *spy-3* mutant. A, C. Representative maps of relative stiffness (arbitrary units) measured on the middle domain of an approximately 150 μm-long growing primordium (rank 11) from WT (A) and *spy*-3 (C) plants grown in SD conditions. Each pixel represents the relative stiffness calculated from a single force-indentation curve. Apico-basal polarity is indicated. Position of tooth (T) as well as sinus (S) are specified. Scale bars are 10 μm. B, D. Projection of relative stiffness (from A,C) on measured leaf topography for the wild type (B) and *spy-3* (D). E. Quantification of relative stiffness cell walls of the tooth (T) or the sinus (S). Each point represents a measured pixel. F. Apparent Young’s modulus (Pa) measured on transversal cell walls of WT teeth (n=11) and sinuses (n=11) and *spy-3* teeth (n=7) and sinuses (n=7).

As CUC2 is required to initiate teeth, it is difficult to use the *cuc2* loss-of-function to assess whether CUC2-dependent inhibition of cell expansion occurs also at the leaf margin. The *spy-3* mutant initiates teeth but their growth is not maintained due to local cellular changes at the sinus during teeth development. *spy-3* is therefore uncoupling CUC2 functions as SPY leads to the repression of the expression of a common set of genes with CUC2 that are related to the cell wall remodeling. We therefore used the *spy-3* mutant to check whether the reduced expression of cell wall genes acting downstream of CUC2 is sufficient to change the mechanical properties of the cell wall at the leaf margin. *spy-3* mutant young leaf primordia show a reduction of the differential relative stiffness between tooth and sinus observed in the WT (Fig 6C-D-E). We confirmed that by quantifying the Apparent Young’s modulus (*Ea*) in another set of experiments analyzing cell wall stiffness in 7 young *spy-3* dissected leaves (Fig 6F). Even though we highlight a significant difference between the sinus and tooth *Ea* values in *spy-3* leaves, this difference was reduced compared with the sinus and tooth *Ea* values of the WT. This data supports the idea that the set of genes commonly down-regulated by CUC2 and SPY may impact cell wall stiffness at the leaf margin. Furthermore, it is important to note that stiffness changes predate the sinus cell morphological changes we observed in the *spy-3* mutant and occur at early stages of leaf development thus providing a mechanical framework for the subsequent differential growth.

## Discussion

Plant development and architecture rely on the iterative production of lateral organs separated from the stem cell pool by boundary domains. Here, through the characterization of leaf margins phenotypes of *spy* mutants at multiple scales, we highlight a role for SPY in maintaining local growth repression of sinus cells redundantly with CUC transcription factors. Further we present evidence that SPY and CUC2 act through a common molecular network involving the reduction of the expression of genes associated with cell wall loosening. Accordingly, using dark-grown hypocotyl together with a CUC2 inducible system, we show that CUC2 restricts cell elongation. This mode of action is supported by the fact that teeth and sinuses of young leaf primordia display differential cell wall stiffness: sinuses - where *CUC2* is expressed - show more rigid cell walls. Accordingly, cell wall stiffness is lower in the sinuses of *spy-3* leaves compared to the WT. Our data support a model where CUC2 can inhibit cell expansion independently of CUC3 acting through the repression of cell wall relaxing genes.

### SPY is a component of the boundary domains network

Here, we have shown that *spy* mutants display boundary domain defects during leaf development resulting in leaves with reduced serrations. Leaf development is a complex and integrated process, which results from both global and local changes throughout developmental time. Therefore we cannot rule out the fact that the local changes we observed in the *spy-3* mutant may results from growth rate changes at the whole organ level. Indeed, GA is involved in the proliferation/differentiation switch that occurs during leaf development. In Arabidopsis, DELLAs were shown to increase the transcript levels of Kip-related protein 2 (KRP2), SIAMESE (SIM) and SIM Related 1 and 2 (SMR1 and SMR2), which are all involved in cell cycle progression inhibition (Achard et al., 2009). As SPY was shown to activate RGA (Zentella et al., 2017), in *spy* mutants, cell cycle progression is less restricted, triggering *in fine* a faster differentiation of the leaf blade. In addition, *GA20ox1* overexpression results in increased levels of bioactive GA and the corresponding lines display large leaves with more and larger cells (Gonzalez et al., 2010). Together these results suggest that GA levels control both cell expansion and cell proliferation in leaves. In addition to its role in GA signaling repression, former studies pointed out that SPY has a role in CK signaling promotion (Greenboim-Wainberg et al., 2005; Steiner et al., 2012). Yet, it was shown that CK promotes cell proliferation and that reduced CK levels lead to a decrease in cell divisions and consequently to smaller organs (Holst et al., 2011). Moreover, a recent study revealed that a CK/GA balance is responsible for leaf complexity in tomato as it controls morphogenesis/differentiation switch (Israeli et al., 2021). Hence, it is also possible that CK signaling is partially impaired in the *spy-3* mutant and as a consequence modifies the serration growth dynamics. Even though we cannot exclude that SPY impacts in multiple ways the global leaf development, we have seen that the *spy-3* mutation results in local changes in cell size, which plaid for a SPY function in boundary definition. Accordingly, SPY acts redundantly with CUC2 and CUC3 to promote cotyledons separation, which is a developmental process where cell expansion mostly occurs in a separated timeframe. It is also important to note that although SPY and CUCs are commonly expressed within leaf boundary domain cells, SPY expression in not restricted to these domains. This observation suggests that sinus-localized defects in the *spy-3* mutant may reflect different activity of SPY depending on the tissue where it is expressed and/or that leaf cells do not respond uniformly to SPY activity alterations.

### SPY and CUC2 act through a common cell wall-related molecular network

At the organ level, the origin of serrations has long been debated (Bilsborough et al., 2011; Kawamura et al., 2010; Nikovics et al., 2006). Recently, morphometrics was used to reconstruct leaf developmental growth trajectories of the wild type and the *cuc2-1* loss-of-function mutant which do not initiate serrations (Biot et al., 2016). This work shows that local growth repression arises first at the sinus of young WT leaf primordia, where CUC2 is expressed, then subsequently, outgrowth appears at a distance. As no growth repression neither teeth initiation were observed in the *cuc2-1* mutant, it was concluded that CUC2 is a key regulator for the coordination of cellular processes leading to serrations development. Our work provides a molecular framework for the growth repression function of CUC2. Indeed, we present evidences that CUC2 and SPY down-regulate a set of common genes involved in cell wall loosening. Accordingly, we demonstrate that CUC2 inhibits cell elongation and that this cellular mode of action is accompanied with down-regulation of transcripts encoding expansin-like and xyloglucan endotransglucosylases/hydrolases which have been reported to be sufficient to promote hypocotyl cell elongation when overexpressed (Boron et al., 2015; Miedes et al., 2013). Unfortunately, no growth phenotypes have been reported for loss-of-function mutants for these genes probably due to the high level of redundancy or the deleterious effects of multiple pleiotropic mutations.

Additionally, we demonstrate here that CUC2 can inhibit cell expansion independently of CUC3. Our current model for CUC2 activity states that *CUC2* is expressed early during leaf development and triggers serration development through CUC3 and KLUH which act as molecular relays (Maugarny-Calès et al., 2019). Accordingly, CUC3 has been shown to inhibit cell expansion of sinus cell hence participating to the shaping of the leaf (Serra and Perrot-rechenmann, 2020). Here, we completed this model by adding another route for the growth repression of sinus cells. As CUC2 can regulate cell expansion independently of CUC3, it is probable that CUC2 contributes also to the local growth repression process *per se*. This reveals an entangled role for CUC2 in coordinating patterning and cell growth to define boundary domains.

### Cell wall mechanics at the leaf margin

Our data show that sinuses and teeth display differential cell wall stiffness even at early stage of leaf development. This differential stiffness is reduced in the *spy-3* mutant where sinuses cell wall resembles more to teeth cell wall. Here, the *spy-3* mutant allows us to decompose CUC2 functions as SPY and CUC2 act on a common molecular network related to cell wall loosening. Our work reveals the contribution of cell wall mechanics to morphogenesis: local cell wall parameters will grandly impinge on the growth of the whole organ. Moreover, CUC3 has been shown to act downstream of CUC2 (Maugarny-Calès et al., 2019), and its expression is promoted by mechanical stresses (Fal et al., 2016). This is therefore tempting to propose a model where CUC2 activity induces mechanical stress at the margins which then could trigger CUC3 expression to serve as a relay for local growth repression. Further experiments will be needed to provide a comprehensive view on the integration of hormonal, genetic and mechanical stress to leaf development and boundary domain development in general.

## Materials and Methods

### Plant material and growth conditions

All plants used in this study were in the *Columbia* (Col-0) background except the *cuc2-1* mutant which was originally obtained in the *Landsberg* (Ler-0) background and back-crossed 5 times to col-0. For morphometric analysis, seeds were immersed in distilled water for two days in the dark at 4°C before sowing on soil. Then, plants were grown under short days conditions (6 hours day [21°C, hygrometry 65%, light 120 μM/m^2^/s], 1 hour dusk [21°C, hygrometry 65%, light 80 μM/m^2^/s], 16 hours night [18°C, hygrometry 65%, dark conditions], 1 hour dawn [19°C, hygrometry 65%, light 80 μM/m^2^/s]. For *in vitro* cultures, seeds were sown on Arabidopsis medium (Gonçalves et al., 2017), stratified for 48 hours in the dark at 4°C then transferred to long day conditions (21°C, 16 hours day / 8 hours night, light 50 μM/m^2^/s).

### Morphometrics analysis

For morphometric analysis of mature leaves, leaves from ranks 11, 12 and 13 from 6-week-old plants were harvested and glued on a paper sheet prior to scanning using a Perfection V800 Photo scanner (Epson) at 1600dpi. For morphometric analysis of developing leaves, young leaf primordia (rank 11, 12 and 13) were dissected using a stereomicroscope throughout development starting at day 22 after sowing. Leaves were mounted between a slide and a coverslip in a buffer containing Tris HCL, 10 mM, pH = 8.5, Triton 0.01% (v/v) and imaged using an Axio Zoom.V16 microscope (Carl Zeiss Microscopy, Jena, Germany; http://www.zeiss.com/). Depending on the developmental stage imaged, either the chlorophyll fluorescent signal or the brightfield signal were collected. Leaf silhouettes and measurements were obtained using the *Morpholeaf* software which allows semi-automatic leaf segmentation and the extraction of relevant biological parameters (Biot et al., 2016). Output data analysis, statistics, and plots were performed using R software (R Core Team, 2016) and the graphics package ggplot2.

### Confocal imaging

For cellular parameters quantification, we used the *pPDF1::mCitrine-KA1* (Stanislas et al., 2018) line in order to visualize the plasma membrane in the leaf epidermis. 26 to 31-day-old Col-0 and *spy-3* plants containing the *pPDF1::mCitrine-KA1* construct were grown under short days conditions prior to dissecting, mounting in a buffer containing Tris HCL, 10mM, pH = 8.5, Triton 0.01% (v/v) and direct imaging with a Leica SP5 inverted microscope (Leica Microsystems, Wetzlar, Germany; http://www.leica-microsystems.com/). Samples were excited using a 514 nm laser and fluorescence was collected with a hybrid detector at between 569 and 611 nm. TIF images were rotated using TransformJ plugin. Subsequent cell segmentations, cell curvature and cell surface area measurements were then obtained using the MorphographX (MGX) (de Reuille et al., 2015) software (http://www.mpipz.mpg.de/MorphoGraphX/). Cells corresponding to tooth sinus were identified as the cells displaying a fully negative signal when projecting a 15 μm-neighboring Gaussian Curvature (Serra and Perrot-rechenmann, 2020). The *pSPY::SPY-GFP, pCUC3::CFP* and *pCUC2::RFP* reporter lines were imaged with a Leica SP5 inverted microscope (Leica Microsystems, Wetzlar, Germany; http://www.leica-microsystems.com/). GFP was excited using a 488 nm laser and fluorescence was collected with a hybrid detector at between 500 and 530 nm. RFP was excited at 561nm and detected with a PMT detector within 570 and 635 nm. CFP was excited at 458nm and detected with a PMT detector within 460 and 475 nm.

### Transcriptomic analysis

#### RNA samples

For RNAseq assays, *p35S:CUC2-GR* seedlings were grown in liquid Arabidopsis medium with constant shaking. After 10 days of growth under constant light, seedlings were treated with DEX or Mock for 6 h and then snap-frozen in liquid nitrogen. DEX (Sigma, D1756) was dissolved in EtOH and used at a final concentration of 10 μM. Total RNA extraction was performed with the miRvana extraction kit (Ambion) following the manufacturer’s recommendations.

#### RNA-seq libraries

RNA-seq libraries were constructed by the POPS platform (IPS2) using the TruSeq no stranded mRNA library prep kit (Illumina®, California, U.S.A.) according to the supplier’s instructions. The libraries were sequenced in paired-end reads (PE, 2×100 bases) on Illumina Hiseq2000 (thanks to IG-CNS to give us a privileged access to perform sequencing) to generate a mean of 30 million of PE reads per sample. Quality process removed PE reads with Qscore < 20, length < 30 bases and ribosomal reads.

#### Bioinformatic analyses

Filtered PE reads were mapped using Bowtie2 (Langmead B., Salzberg SL. 2012) with the --local option against the *Arabidopsis thaliana* transcriptome. The 33602 genes were extracted from TAIR v10 database. 95% of PE reads were associated to gene without ambiguously, 2% removed for multi-hits. Genes with less than 1 read per million in at least half of the samples were discarded. The resulting raw count matrix was fed into edgeR (Robinson et al., 2010) for differential expression testing by fitting a negative binomial generalized log-linear model (GLM) including a condition factor and a replicate factor to the TMM-normalized read counts for each gene. We performed pairwise comparisons of each of the DEX-treated condition to the control condition. The distribution of the resulting p-values followed the quality criterion described by Rigaill et al. 2018. Genes with an adjusted p-value (FDR, Benjamini-Hochberg adjustment) below 0.05 were considered as differentially expressed.

#### Data Availability

RNA-Seq projects were sent to GEO/NCBI (Edgard R. et al. 2002): http://www.ncbi.nlm.nih.gov/geo/; accession no. GSE184530. All steps of the experiment, from growth conditions to bioinformatic analyses, were detailed in POPS database, CATdb (Gagnot S. et al. 2007): http://tools.ips2.u-psud.fr.fr/CATdb/; Project NGS2014_39_LeafNet.

### Dark grown hypocotyl measurements

For dark grown hypocotyls experiments, seeds were surface sterilized and subsequently dispatched on 1% agar (w/v) Arabidopsis media with DEX (Sigma, D1756) at 10 μM or Mock treatment. After 48 hours in the dark at 4°C, plates were transferred to growth chamber for 6 hours (21°C, light 50μM/m^2^/s) before being placed vertically in the dark at 21°C. Plates were scanned after 72 hours of dark growth using a Perfection V800 Photo scanner (Epson) at 1600 dpi. Hypocotyl sizes were measured using the NeuronJ plugin from Fiji and data were analyzed using R software (R Core Team, 2016).

### Expression data

Total RNA were isolated using RNAeasy Plant Mini Kit (Qiagen) following the manufacturer recommendation for plant tissue. Reverse transcription was performed using RevertAid H Minus M-MuLV Reverse transcriptase (Fermentas) followed by a RNAse H treatment was performed for 20 min 37°C to eliminate DNA-RNA duplexes. Real time PCR analysis was performed on a 384-well QuantStudio™ 5 Real-Time PCR System, using the SsoAdvance Universal SYBR Green Supermix with the following PCR conditions: 95°C 3min; (95°C 15 s; 63°C 30 s) x 40 cycles. Raw data was analyzed using Design & Analysis 2.2 software. Primers used for real time PCR analysis are available in Table S8. Expression data were normalized using the ΔΔCt method using at least two independent reference genes.

### Atomic Force Microscopy (AFM)

For mechanical characterization of leaf cell wall, WT and *spy-3* plants were grown on soil in short-day conditions. About 100-200 μm-long young leaf primodia from rank higher than 11 were hand-dissected under a stereomicroscope and collected. They were fixed in low melting agarose blade facing the AFM tip following the protocol described in (Peaucelle, 2014). Samples were plasmolysed by immersion in sorbitol 10% (m/v). An AFM cantilever loaded with a 1μm diameter tip was used in these measurements and scanned 100 μm*30 μm areas with a fixed force leading to a maximum indentation value of 800 nm with a speed of 40 μms^-1^. The measurement of the rigidity constant was performed only on the second cantiliever used as a reference tip as described in (Peaucelle, 2014). Apparent Young’s modulus was determined by a Hertzian indentation model on each indentation point. Cells topography was reconstructed using the height at each point of contact. Data were analyzed and maps were plotted using Matlab.

## Acknowledgments

We thank Utpal Nath (Indian Institute of Science Bangalore) for the gift of *spy-3* seeds, Doris Lucyshyn (BOKU, Vienna, Austria) for sharing with us the *spy-22* and *spy-23* mutants, the pSPY::SPY-FLAG *spy-22* and the pSPY::SPY-GFP lines and Ben Scheres (Wageningen University, the Netherlands) for the gift of the p35S:CUC2-GR line. We thank Véronique Pautot and Antoine Nicolas for discussions and critical reading of the manuscript. This work was supported by the Agence National de la Recherche grants LEAFNET (ANR-12-PDOC-0003) and by the INRAE grant MorphEAT (AAP BAP 2020). The IJPB benefits from the support of the Labex Saclay Plant Sciences-SPS (ANR-10-LABX-0040-SPS).

## Author Contributions

NB, PL and NA: designed the research; PL and NA acquired funding, NB, AP, MG, BA, and NA performed research; LST, ZT and MLMM performed transcriptomic analysis; NB and AP performed AFM experiments; EB contributed to new computational tools; NB, AP and NA analyzed data; NB and NA wrote the paper with contributions from PL.

